# The unintended consequences of a surge in antibiotic resistance when removing wastewater drain biofilm

**DOI:** 10.1101/2020.11.06.372326

**Authors:** Shireen M Kotay, Hardik I Parikh, Hyun S Gweon, Katie Barry, Nicole Stoesser, A Sarah Walker, Derrick W Crook, Kasi Vegesana, Amy J Mathers

## Abstract

The prevalence and proliferation of antimicrobial resistant bacteria is considered one of the critical issues of our time. Wastewater is a habitat for complex microbial communities where bacteria share genes of antimicrobial resistance through horizontal gene transfer. Hospital wastewater drainage systems are an ideal reservoir for environmental and pathogenic bacteria to interface and exchange genes of antimicrobial resistance^1^. Highly resistant Enterobacterales are of the greatest concern because they not only cause human infections but can also thrive in the environment^2,3^ and have shown to transmit from sink drains to patients^4^. Replacement of contaminated plumbing may be the most intuitive and widely deployed response once highly resistant potentially pathogenic bacteria are identified in a sink drain. Here we show using shotgun metagenomic sequencing of sink drain biofilms (in 6 intensive care patient rooms, 36 subsamples) paired with microbial culture analysis, an evident shift of biofilm community structure towards increased abundance of Enterobacteriaceae in the newly replaced sink plumbing. Significant increase in resistome load and abundance of clinically relevant resistance and typically mobile genes in the newly replaced plumbing was also observed. Findings from this study demonstrate that exchanging contaminated plumbing for new plumbing may actually have the unexpected consequence of increased abundance of Enterobacterales and genes of antimicrobial resistance in the sink drains. Disruption of preexisting complex environmental biofilms may result in an unintended increase in the amount of antimicrobial resistant Enterobacterales which are targeted for elimination. Temporal variability is an important attribute of ecosystems affecting colonization dynamics and biotic interactions (composition of introduced and resident communities) are key to functional outcomes. Absence of competition within an ecosystem favors invasion.

## MAIN

Antimicrobial resistance has emerged as a major threat to human health, according to a United Nations 2019 report^5^. Some of the most concerning antimicrobial resistance is driven by the ability of pathogenic bacteria to exchange antimicrobial resistance genes. Enterobacterales which can acquire carbapenem resistance genes via horizontal gene transfer are among the most clinically relevant antimicrobial resistant bacteria, with associated infections resulting in high morbidity and mortality^6^. Wastewater has been identified as a domain which can facilitate mixing of diverse bacteria enabling the exchange of some of the most consequential genes of antimicrobial resistance^7^. This is particularly influenced by compounds which may be present in wastewater such as antibiotics, disinfectants, antimicrobials and heavy metals, which can exert selection pressures even in low concentrations or in their derivative forms.

The microbiology of the built environment is heavily influenced by the use and inhabitants of that environment^8^. Hospitals represent a habitat harboring some of the most resistant bacteria carried by patients with heavy antimicrobial usage and a high proportion of human pathogens compared to other built environments^9^. Further, highly antibiotic resistant pathogens and opportunistic pathogens are now frequently detected in the wastewater plumbing of hospital sinks and toilets with reports of transmission to vulnerable patients^2^. Environmental reservoirs such as sinks in patient care areas contaminated with highly resistant bacteria are frequently targeted for mitigation under the auspices of patient safety. Intervention measures have included replacement of plumbing^10–15^, and/or the use of chemical disinfectants such as bleach^12^, acetic acid^13^, and hydrogen peroxide^11^. Plumbing replacement is a widely implemented intervention strategy and was found to be a relatively more effective method in mitigating outbreaks^3^. However, duration of follow-up after many intervention studies has been short, and the longer-term impact on plumbing-associated bacterial communities remains unevaluated^12-14,16-19^.

Having experienced a prolonged outbreak of carbapenemase-producing Enterobacterales (CPE) associated with wastewater plumbing contamination and persistence in our institution, we set out to monitor the dynamics around removal of sink plumbing and recolonization of the *in situ* biofilm in newly replaced sink drains. We studied six intensive care room hand wash sinks, all of which had similar design. Sink drains were sampled at monthly intervals for three months before and after plumbing exchange. Bacterial culture and shotgun metagenomics of the collected drain biofilm was performed^20^. The culture data depict the increase in the proportion of sampling events where sink drains were positive for multiple species of CPE (Fig 1a bottom panel); 39% (7/18) before as compared to 89% (16/18) after plumbing replacement (p= 0.006). In addition, the number of CPE isolates per sampling occasion increased from 0.83 (15/18) to 2 (36/18) pre- and post-plumbing exchange, respectively. Two of the six sinks (Rooms 1 and 3) that were culture-negative at all sampling timepoints for three months prior to plumbing replacement, were first detected to be positive for CPE after plumbing replacement. In all but one room (Room 1), CPE isolates were cultured immediately (sampling timepoint-4) after plumbing replacement and were persistently detected in the following months.

**Figure 1:**
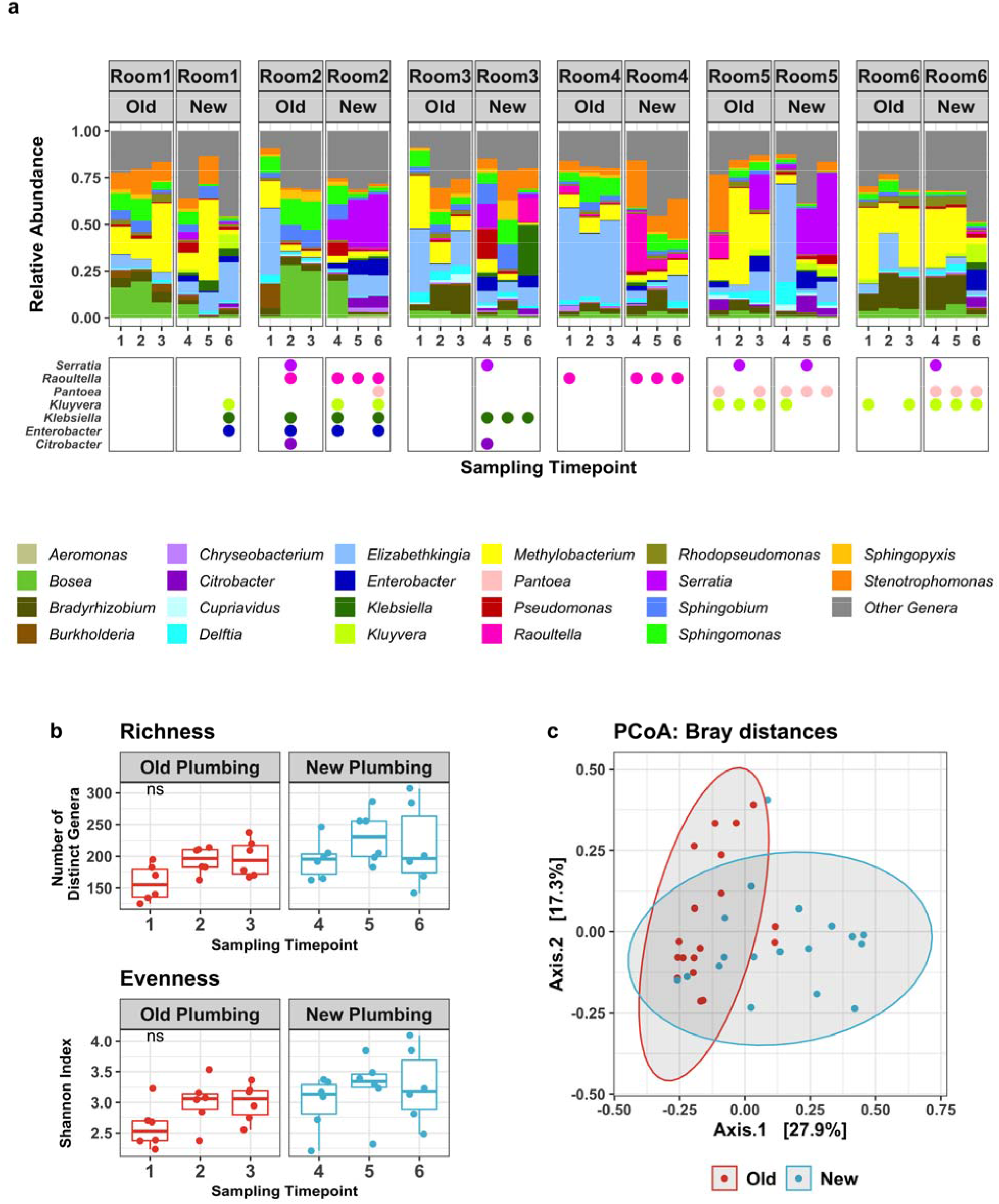
Microbial community composition in sink microbiomes. a) Top panel: Relative abundance of bacterial genera in the sink microbiomes grouped by old (time points 1, 2, 3) and new plumbing (time points 4, 5, 6) for each room. Abundances of select genera are shown as stacked bars, with remaining low abundance genera aggregated into *Other.* Bottom panel: Cultured CPE isolates from sink drain biofilms. b) Boxplots showing alpha diversity of the microbial community, represented as richness and evenness (Shannon index), stratified by old plumbing (time points 1, 2, 3) and new plumbing (time points 4, 5, 6). The horizontal box lines represent the median and the inter-quartile range. No evidence of change observed over time (Kruskal-Wallis; NS; p-value >0.05). c) Beta diversity at genus-level among microbial communities of sink metagenomes, explored by principal coordinate analysis of pairwise Bray-Curtis dissimilarity indices.

Metagenomic taxonomic diversity analysis revealed a largely dissimilar and variable community composition and structure across sinks, samples and timepoints (Fig. 1a top panel). Replacing the plumbing showed no significant changes in the community richness and evenness at genus-level (Fig. 1b and c). This is in contrast to the shift in microbiome diversity seen in the setting of disinfection of hospital surfaces where there was a decrease in diversity^9^. Despite no significant changes in the overall diversity, there was a significant increase in the Enterobacterales population post-plumbing replacement including genera commonly associated with nosocomial infections and antimicrobial resistance such as *Serratia, Enterobacter, Klebsiella* and *Citrobacter* (Fig. 2, Fig. S1). In contrast, the relative abundance of slow-growing, native dwellers of water environments like *Methylobacterium, Bosea* and *Sphingomonas* spp. were negatively associated with plumbing exchange.

**Figure 2.**
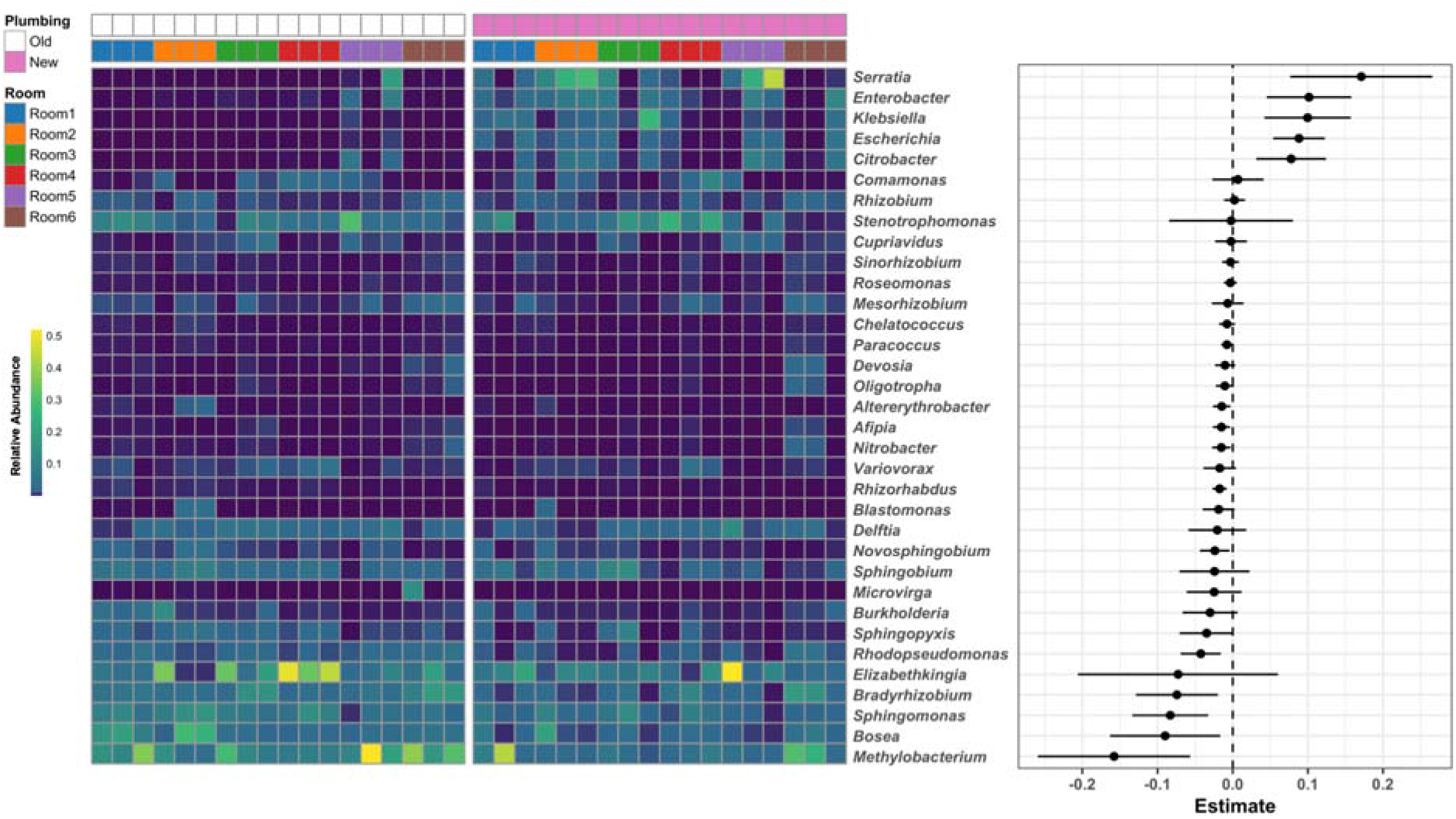
Relative abundance (left panel; heat map) and estimates using linear mixed effects modeling (right-panel; points indicate estimate values, with line showing 95% CI) for significantly differentially abundant genera in new sink plumbing compared to old plumbing samples. Samples are grouped by old and new plumbing, with samples collected from same room further grouped together (as indicated by annotation bars at top). (See Supplementary Table S2 for estimates, 95%CI, p-values and q-values for each taxa).

The shift in bacterial community structure towards Enterobacterales after plumbing replacement was positively associated with the significant increase in total resistance gene load (Fig. 3). Relative abundances of genes conferring resistance to beta-lactams (*bla*_KPC-2_, *bla*_OXA-32_ and *bla*_TEM-3_ clusters) and aminoglycosides that are mostly plasmid-borne and frequently found in CPE, were significantly higher in the new plumbing (Fig. 4). As shown by the pairwise analysis (Fig. S2), relative abundance of these genes and Enterobacterales were strongly correlated. New plumbing with noticeably higher *bla*_KPC_ abundance corresponded with a remarkably higher abundance of *Klebsiella, Enterobacter* and *Serratia.* Isolates belonging to these genera frequently carrying the carbapenemase gene, *bla*_KPC_, have been isolated from patients and plumbing in this setting^20^; and all the CPE identified from culturing plumbing biofilm in this study were *bla*_KPC_.

**Figure 3.**
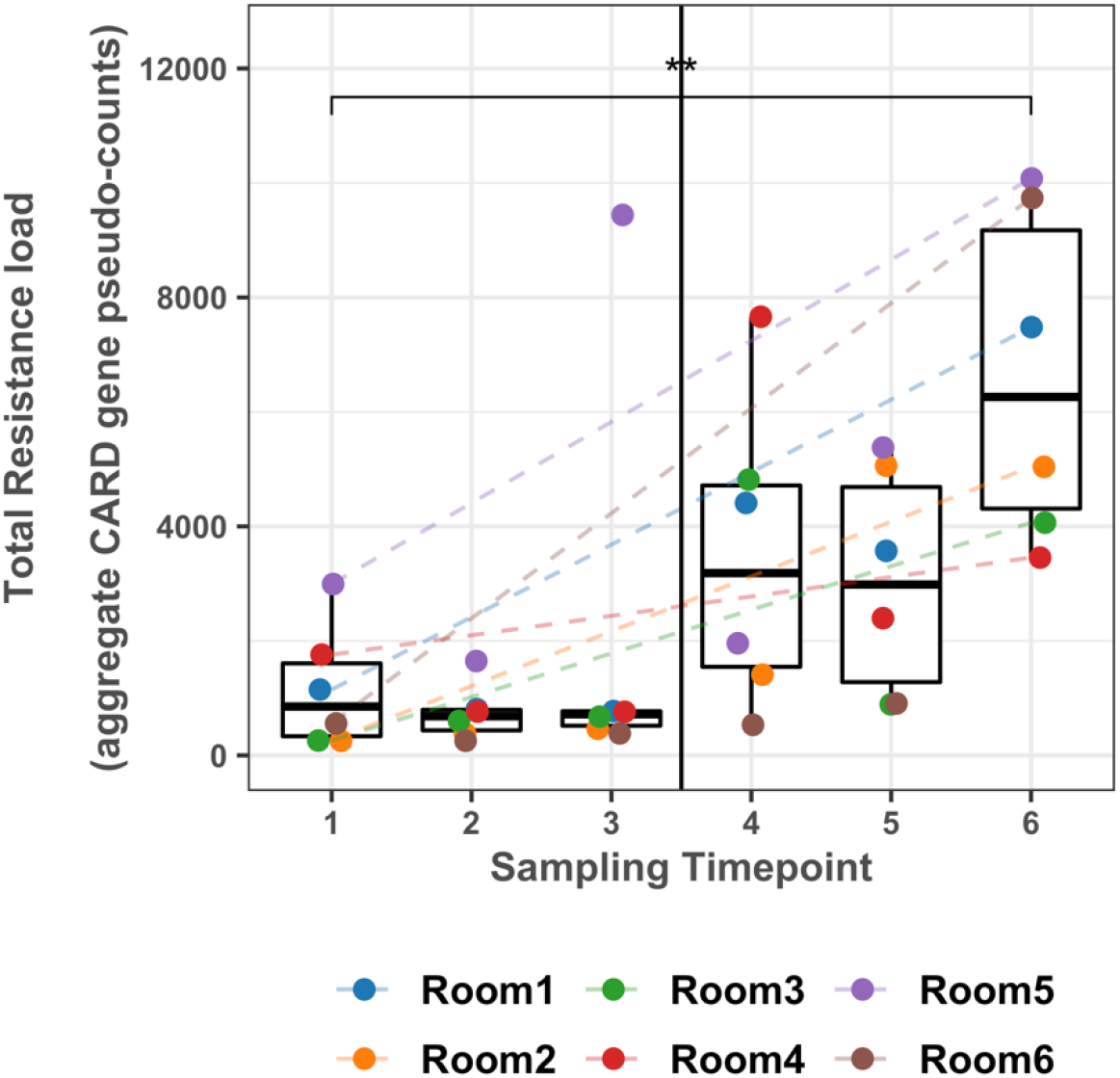
Box plots showing total sum abundances of resistance genes (aggregate pseudo-counts; see Methods for details) in sink biofilms; stratified by old plumbing (time points 1, 2, 3) and new plumbing (time points 4, 5, 6). The horizontal box lines represent the median and the inter-quartile range. There is a significant increase in total resistome load in old plumbing samples collected at time point 1 compared to new plumbing samples collected at time point 6 (two-sided paired Wilcoxon test; n=12; p=0.002).

**Figure 4.**
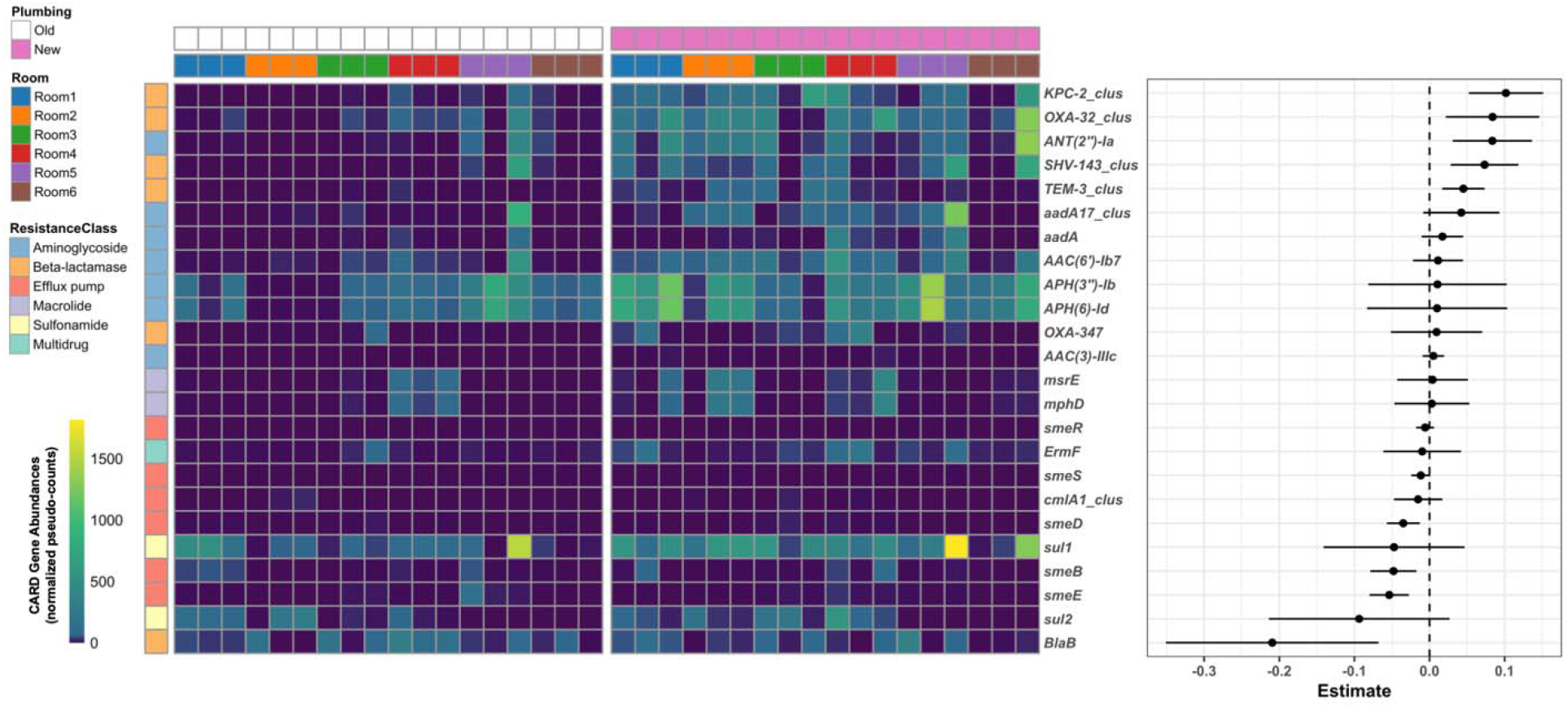
Relative abundance (left panel; heat map) and estimates using linear mixed effects modeling, (right-panel; points indicate estimate values, with line showing 95% CI) for significantly differentially abundant antimicrobial resistance genes in new sink plumbing compared to old plumbing samples. Samples are grouped by old and new plumbing, with samples collected from same room further grouped together (as indicated by annotation bars at top). The annotation bar to the left indicate the resistance class for resistance gene. (See Supplementary Table S2 for estimates, 95%CI, p-values and q-values for each gene).

To investigate the potential effect of external factors, we compared the room attributes and patient factors that might have affected the microbial community in the sink drain biofilms. Whilst power was limited, there were no statistically significant differences in factors that directly or indirectly impact the sink usage in the rooms before and after plumbing replacement (Table S1).

Studies have demonstrated rapid recolonization with drug-resistant potential pathogens after hospital plumbing exchange,^13,14,21^ and this systematic study supports those findings, but highlights in much greater detail the associated wider microbiome shifts that occur over short timeframes when sink drains are replaced. The rapid recolonization of newly replaced plumbing by Enterobacterales is analogous to an invasive event, where a disrupted natural population may be replaced by a less desirable invasive population. Although it might make intuitive sense to exchange drain plumbing when antimicrobial-resistant pathogens are identified, this may have the unintended consequence of destabilizing a microbial community, disrupting colonization resistance and thus promoting the successful overgrowth of the antimicrobial resistant pathogens which were originally targeted for removal. Rather, mitigation strategies and interventions should potentially focus on keeping the drain free from pathogen colonization and on barriers that can prevent dispersal from sinks to patients. Altering the structure of the sink drain microbiome alters the function of the community, which likely reflect changes in the ecologic, resistance, and metabolic functions of the microbiome and thereby facilitate colonization with non-native potentially more drug-resistant, pathogenic communities.

## METHODS

### Sample Collection and processing

Sink drain biofilms were prospectively collected at a monthly interval from six patient rooms in medical and neurosurgical intensive care units for three months before and after the plumbing fixtures (drain, tailpipe, P-trap, trap-arm) were replaced. Pre-sterilized wooden applicator sticks (SKU#: 807, Puritan Medical Products LLC., Guilford, ME) were inserted 1 inch below the sink drain and swirled on the luminal surface to collect the biofilm samples. Collected biofilm was re-suspended into 1ml saline solution and vortexed thoroughly to homogenize. Aliquots of homogenized biofilm suspension were cultured to quantify carbapenem-resistant organisms by directly plating on Colorex KPC agar (North East Laboratories, Waterville, Maine), as well as after enrichment in tryptic soy broth containing 10μg ertapenem disc, and subsequent plating on Colorex KPC agar. Species identification of cultured isolates was performed using the VITEK2 (Biomerieux, Durham, NC); isolates were also screened for *bla*_KPC_ by PCR as previously described^20^. Total DNA was extracted from the biofilm samples using the PowerSoil DNA Isolation kit (QIAGEN) following the manufacturer’s protocol. After quantification using picogreen the extracted DNA was used to construct shotgun metagenomic libraries using the Nextera XT kit following the manufacturers’ protocols. Sequencing was performed using the Illumina HiSeq 4000, multiplexing six samples per lane to generate approximately 40 million 150 paired-end reads per sample.

### Metagenomic analyses

Low-quality portions of sequences were trimmed using TrimGalore^22^. Specifically, we discarded reads below an average Phred score of Q25 and a length cut-off of 75bp, and the first 13bp of Illumina adapters. Estimation of taxonomic composition of all metagenomics samples was performed with Kraken^23^ using NCBI’s RefSeq bacterial and viral genomes as reference database (accessed: 02/2018). For each taxonomic level, relative abundances were estimated using Bracken^24^. To remove spurious and low-abundance assignments, only genera with >0.01% relative abundance was considered for downstream diversity and statistical analysis.

For AMR gene abundance estimation, the filtered metagenomic reads were mapped using bbmap^25^ against the Comprehensive Antibiotic Resistance Database (CARD, v2.0.1)^26^. As short reads do not provide sufficient resolution to distinguish highly similar AMR gene homologs, we pre-clustered the CARD database (n=2,448 sequences) at 90% identity and only retained one representative sequence for each cluster (n=1,074 sequences). The aligned BAM files were used for gene counting, normalization and abundance estimation using the approach implemented within ResPipe^27^. Briefly, gene cluster counts were generated for reads that mapped with 100% sequence identity to representative sequence, followed by pseudo-abundance estimation by normalizing using the gene length as well as lateral coverage of the gene sequence by reads.

### Statistical analyses

Within-sample richness and evenness (alpha diversity) of microbial communities were determined by computing the species count and Shannon Index, respectively, using the *vegan* package (v2.5-6)^28^ in R (v3.5.2). Pairwise compositional dissimilarity (beta diversity) was quantified using the Bray-Curtis index. To visualize the dissimilarity between samples, we employed Principal Coordinates Analysis on Bray distances, implemented within the *phyloseq* package in R. For differential abundance analysis, genus-level taxonomic relative abundance profiles were converted to arcsine square-root transformed proportions and random effects model estimates were calculated, using room as a random effect to account for within-room heterogeneity. P-values obtained from random effects models were corrected for multiple comparisons by computing the False Discovery Rate, with resulting q-values <0.1 considered statistically significant. Similarly, resistance gene counts were analysed using random effects models directly on the pseudo-counts. Hierarchical all-against-all association testing was performed to find grouped linear associations between genus-level abundances and AMR gene abundances, using *HALLA*^29^.

### Room attributes collection

For each environmental sampling event, attributes from all six rooms and data on antibiotic usage was gathered from our health system data warehouse (Table S1). Attributes that could potentially have an influence on sink usage and in turn on drain biofilm microbial community (CRE positive patients and antibiotic selective pressure) were included. Data from the three months before and after plumbing replacement for each room were aggregated.

## Data availability

The metagenomic sequences are available at NCBI’s Sequence Read Archive (SRA) under the BioProject ID: PRJNA587635.

---

**Figure S1.**
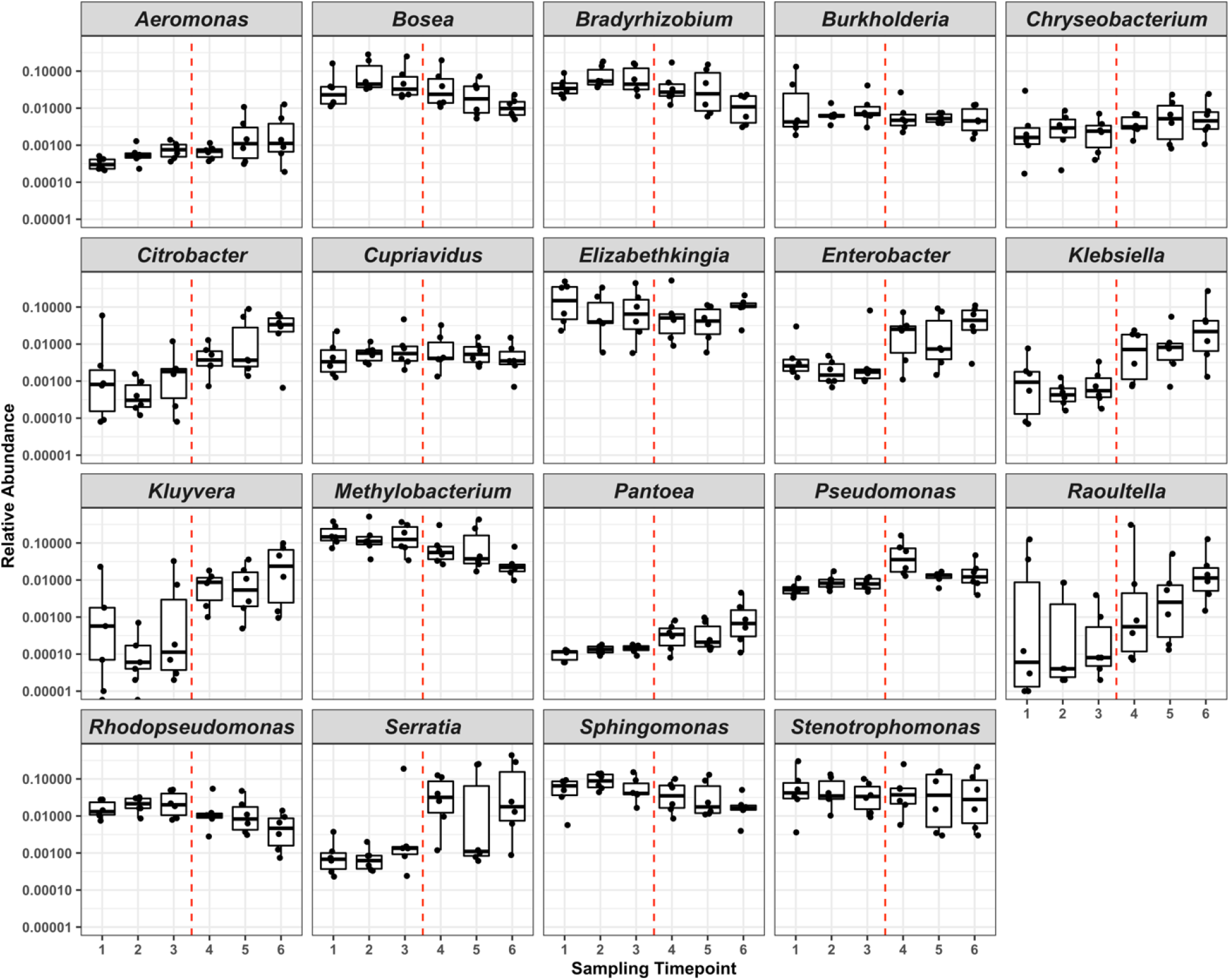
Boxplots showing longitudinal changes in relative abundance of select genera in sink biofilms, stratified by old plumbing (time points 1, 2, 3) and new plumbing (4, 5, 6) indicated by a dashed-red line. The horizontal box lines represent the median and the inter-quartile range. (Note: Y-axis has been log-transformed to visualize changes in very low abundance taxa).

**Figure S2.**
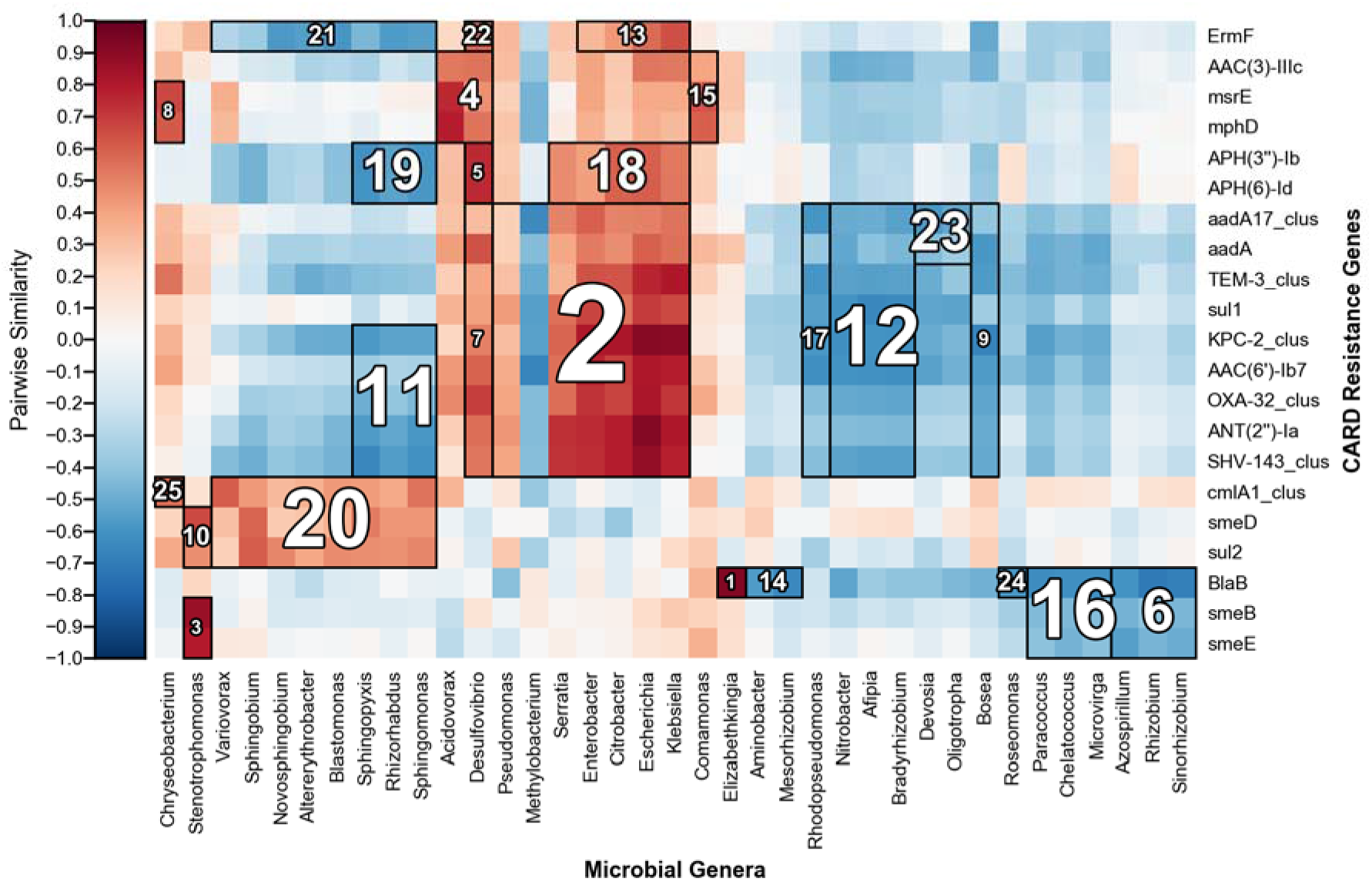
A multi-resolution association plot (HAllA tool^29^, huttenhower.sph.harvard.edu/halla) showing grouped associations in pairwise Spearman’s correlations between genus-abundances and AMR gene abundances in sink biofilms. The number within each group represents association rank.

**Supplemental Table S1.**
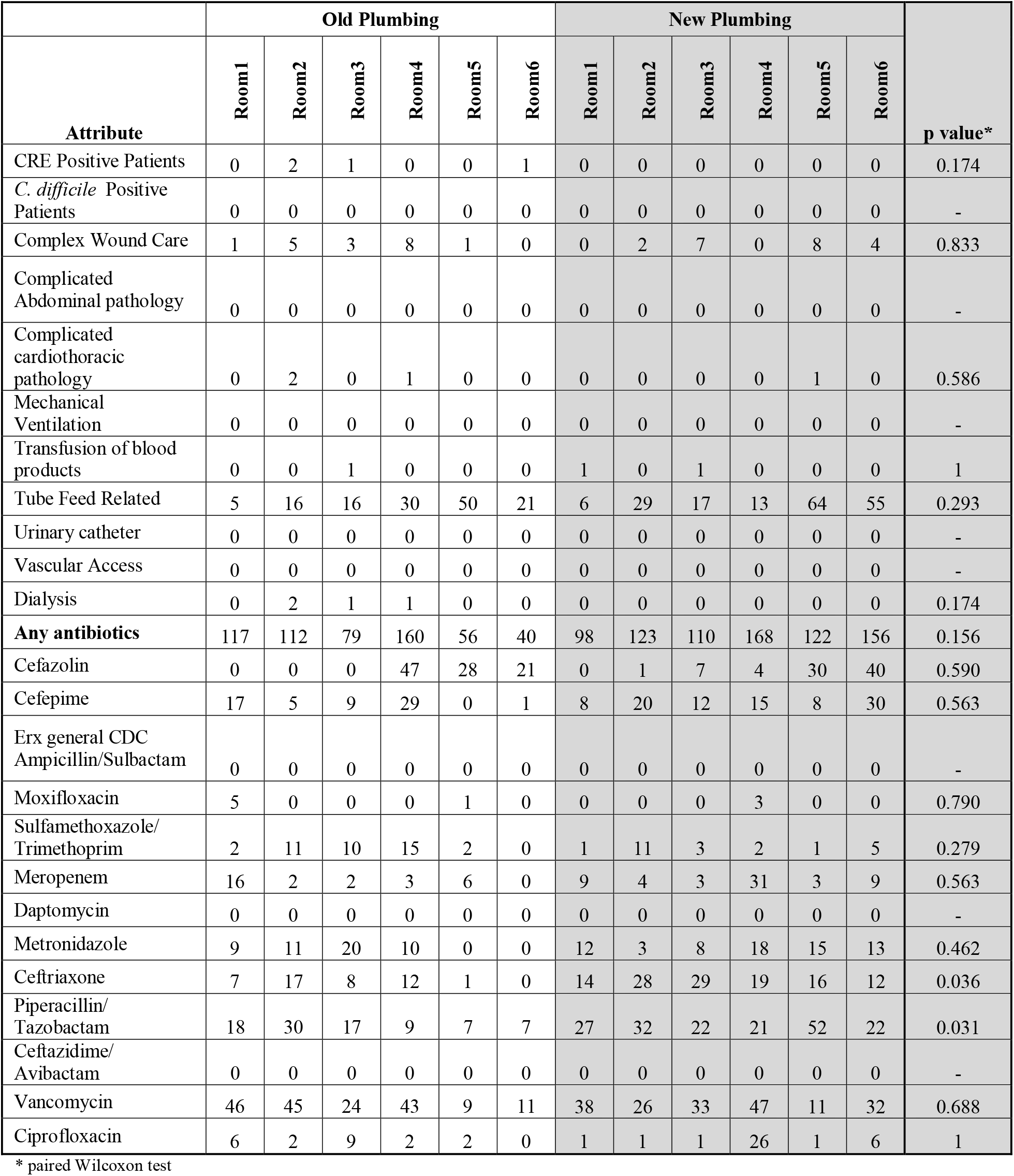
Room attributes represented as cumulative patient days 3months before (Old plumbing) and after (New plumbing) plumbing exchange.

|  | Old Plumbing |  |  |  |  |  | New Plumbing |  |  |  |  |  | p value* |
| --- | --- | --- | --- | --- | --- | --- | --- | --- | --- | --- | --- | --- | --- |
| Attribute | Room1 | Room2 | Room3 | Room4 | Room5 | Room6 | Room1 | Room2 | Room3 | Room4 | Room5 | Room6 |  |
| CRE Positive Patients | 0 | 2 | 1 | 0 | 0 | 1 | 0 | 0 | 0 | 0 | 0 | 0 | 0.174 |
| *C. difficile* Positive Patients | 0 | 0 | 0 | 0 | 0 | 0 | 0 | 0 | 0 | 0 | 0 | 0 | - |
| Complex Wound Care | 1 | 5 | 3 | 8 | 1 | 0 | 0 | 2 | 7 | 0 | 8 | 4 | 0.833 |
| Complicated Abdominal pathology | 0 | 0 | 0 | 0 | 0 | 0 | 0 | 0 | 0 | 0 | 0 | 0 | - |
| Complicated cardiothoracic pathology | 0 | 2 | 0 | 1 | 0 | 0 | 0 | 0 | 0 | 0 | 1 | 0 | 0.586 |
| Mechanical Ventilation | 0 | 0 | 0 | 0 | 0 | 0 | 0 | 0 | 0 | 0 | 0 | 0 | - |
| Transfusion of blood products | 0 | 0 | 1 | 0 | 0 | 0 | 1 | 0 | 1 | 0 | 0 | 0 | 1 |
| Tube Feed Related | 5 | 16 | 16 | 30 | 50 | 21 | 6 | 29 | 17 | 13 | 64 | 55 | 0.293 |
| Urinary catheter | 0 | 0 | 0 | 0 | 0 | 0 | 0 | 0 | 0 | 0 | 0 | 0 | - |
| Vascular Access | 0 | 0 | 0 | 0 | 0 | 0 | 0 | 0 | 0 | 0 | 0 | 0 | - |
| Dialysis | 0 | 2 | 1 | 1 | 0 | 0 | 0 | 0 | 0 | 0 | 0 | 0 | 0.174 |
| **Any antibiotics** | 117 | 112 | 79 | 160 | 56 | 40 | 98 | 123 | 110 | 168 | 122 | 156 | 0.156 |
| Cefazolin | 0 | 0 | 0 | 47 | 28 | 21 | 0 | 1 | 7 | 4 | 30 | 40 | 0.590 |
| Cefepime | 17 | 5 | 9 | 29 | 0 | 1 | 8 | 20 | 12 | 15 | 8 | 30 | 0.563 |
| Erx general CDC Ampicillin/Sulbactam | 0 | 0 | 0 | 0 | 0 | 0 | 0 | 0 | 0 | 0 | 0 | 0 | - |
| Moxifloxacin | 5 | 0 | 0 | 0 | 1 | 0 | 0 | 0 | 0 | 3 | 0 | 0 | 0.790 |
| Sulfamethoxazole/Trimethoprim | 2 | 11 | 10 | 15 | 2 | 0 | 1 | 11 | 3 | 2 | 1 | 5 | 0.279 |
| Meropenem | 16 | 2 | 2 | 3 | 6 | 0 | 9 | 4 | 3 | 31 | 3 | 9 | 0.563 |
| Daptomycin | 0 | 0 | 0 | 0 | 0 | 0 | 0 | 0 | 0 | 0 | 0 | 0 | - |
| Metronidazole | 9 | 11 | 20 | 10 | 0 | 0 | 12 | 3 | 8 | 18 | 15 | 13 | 0.462 |
| Ceftriaxone | 7 | 17 | 8 | 12 | 1 | 0 | 14 | 28 | 29 | 19 | 16 | 12 | 0.036 |
| Piperacillin/Tazobactam | 18 | 30 | 17 | 9 | 7 | 7 | 27 | 32 | 22 | 21 | 52 | 22 | 0.031 |
| Ceftazidime/Avibactam | 0 | 0 | 0 | 0 | 0 | 0 | 0 | 0 | 0 | 0 | 0 | 0 | - |
| Vancomycin | 46 | 45 | 24 | 43 | 9 | 11 | 38 | 26 | 33 | 47 | 11 | 32 | 0.688 |
| Ciprofloxacin | 6 | 2 | 9 | 2 | 2 | 0 | 1 | 1 | 1 | 26 | 1 | 6 | 1 |

\* paired Wilcoxon test

